# Lestaurtinib’s antineoplastic activity converges on JAK/STAT signaling to inhibit advanced forms of therapy resistant ovarian cancer

**DOI:** 10.1101/2024.06.06.597753

**Authors:** Esther P.B. Rodman, Michael J. Emch, Xiaonan Hou, Archit Bajaj, Nicole A. Pearson, August J. John, Yamillie Ortiz, Adam D. Bass, Saloni Singh, Gustavo Baldassarre, Scott H. Kaufmann, S. John Weroha, John R. Hawse

## Abstract

Ovarian cancer is the deadliest gynecological malignancy, owing to its late-stage diagnosis and high rates of recurrence and resistance following standard-of-care treatment, highlighting the need for novel treatment approaches. Through an unbiased drug screen, we identified the kinase inhibitor, lestaurtinib, as a potent antineoplastic agent for chemotherapy- and PARP-inhibitor (PARPi)-sensitive and -resistant ovarian cancer cells and patient derived xenografts (PDXs). RNA-sequencing revealed that lestaurtinib potently suppressed JAK/STAT signaling and lestaurtinib efficacy was shown to be directly related to JAK/STAT pathway activity in cell lines and PDX models. Most ovarian cancer cells exhibited constitutive JAK/STAT pathway activation and genetic loss of STAT1 and STAT3 resulted in growth inhibition. Lestaurtinib also displayed synergy when combined with cisplatin and olaparib, including in a model of PARPi resistance. In contrast, the most well-known JAK/STAT inhibitor, ruxolitinib, lacked antineoplastic activity against all ovarian cancer cell lines and PDX models tested. This divergent behavior was reflected in the ability of lestaurtinib to block both Y701/705 and S727 phosphorylation of STAT1 and STAT3, whereas ruxolitinib failed to block S727. Consistent with these findings, lestaurtinib additionally inhibited JNK and ERK activity, leading to more complete suppression of STAT phosphorylation. Concordantly, combinatorial treatment with ruxolitinib and a JNK or ERK inhibitor resulted in synergistic antineoplastic effects at dose levels where single agents were ineffective. Taken together, these findings indicate that lestaurtinib, and other treatments that converge on JAK/STAT signaling, are worthy of further pre-clinical and clinical exploration for the treatment of highly aggressive and advanced forms of ovarian cancer.

**Statement of significance:** Lestaurtinib is a novel inhibitor of ovarian cancer, including chemotherapy- and PARPi-resistant models, that acts through robust inhibition of the JAK/STAT pathway and synergizes with standard-of-care agents at clinically relevant concentrations.

## Introduction

Epithelial ovarian cancer is the 6th leading cause of cancer related deaths in women and the most fatal of all female reproductive cancers^1,2^. Patient outcomes following the diagnosis of ovarian cancer are dismal with 5-year overall survival of less than 30%^2^. These poor survival rates reflect an insufficiency of routine screening methods, lack of early detection, and *de novo* or rapidly acquired resistance to current standard-of-care regimens^3^. Problematically, over 75% of patients are diagnosed with stage III or IV disease, where there is already metastatic spread throughout the peritoneum with the involvement of other organ sites^3^. Current treatment guidelines for patients with newly diagnosed advanced ovarian cancer includes debulking surgery with adjuvant or neoadjuvant therapy with carboplatin and paclitaxel^4,5^. More recently, a few targeted therapies have been FDA approved, including multiple PARP inhibitors (PARPi)^6^, the anti-VEGF inhibitor, bevacizumab, for use in combination with chemotherapy or a PARPi^7^, and mirvetuximab soravtansine, an antibody-drug conjugate targeting the folate receptor alpha (FRα)^8^. Despite these advances, the majority of patients experience only modest increases in progression-free survival (PFS) with PARPi or VEGFi therapies^6^, and mirvetuximab soravtansine is currently only approved for patients who have received 1-3 lines of prior systemic therapy^8^. Importantly, none of these additional lines of treatment have led to meaningful increases in overall survival.

To combat these challenges, an increasing number of studies have aimed to define mechanisms of ovarian cancer resistance, particularly to platinum-based and PARPi therapies, and to characterize the molecular alterations developed during treatment that may be therapeutically actionable. Thus far, identified mechanisms of resistance include restoration of homologous recombination (HR) activity, increased activity of drug efflux pumps, mitigation of replication stress and alterations in growth factor signaling pathways, among others^9^. Despite these efforts and the many resulting early phase clinical trials, the 5- and 10-year relative survival rates of ovarian cancer patients has only improved by 6.1% and 2.5% respectively since the 1980’s^10^. These realities highlight the need to better understand the basis for systemic treatment failure and to devise alternative, more effective treatment strategies.

In this study, we employed an unbiased drug screen to identify small molecules that potently inhibit and/or kill both treatment-sensitive and treatment-resistant ovarian cancer cell line models. We identified the kinase inhibitor lestaurtinib as a drug of interest and, through RNA-sequencing (RNAseq) analysis, demonstrated that this agent primarily inhibits the JAK/STAT signaling pathway in ovarian cancer cells. Further analysis comparing lestaurtinib to the FDA approved JAK inhibitor, ruxolitinib, found that lestaurtinib inhibited STAT1 and STAT3 serine as well as tyrosine phosphorylation. Finally, lestaurtinib showed monotherapy activity in many ovarian cancer cell line and PDX models and enhanced the antineoplastic effects of cisplatin and olaparib in both treatment naïve and treatment refractory models.

## Methods

### Drugs and recombinant proteins

The epigenetics screening library (#11076) and lestaurtinib (#12904) were purchased from Cayman Chemicals. Ruxolitinib (#S1378), olaparib (#S1060), and LY3214996 (#S8534) were purchased from SellekChem. CC-90001 (#HY-138304) was purchased from MedChemExpress. Cisplatin was purchased from Fresenius Kabi (#100351). Recombinant proteins, IFNγ (#285-IF) and IL6 (#7270-IL), were purchased from R&D Systems.

### Cell culture

MDAH-2774 (MDAH) and OVSAHO cells, and their respective platinum-resistant clones^11^, were generously provided by Dr. Gustavo Baldassarre (National Cancer Institute, Italy). PEO1 and PEO4 cells were derived from the same patient during early and later points of her care^12^, and were generously provided by Dr. Scott Kaufmann (Mayo Clinic, Rochester, MN). The ABT^R^#2 cell line was derived from PEO1 cells by PARPi selection and were characterized as previously described^13^, COV362 cells and an olaparib-resistant derivative COV362-Olap^R^ generated from COV362 by continuous selection in olaparib concentrations from 0.5 µM to 10 µM (characterized as illustrated in Supplemental Figure 1), were generously provided by Dr. Scott Kaufmann (Mayo Clinic, Rochester, MN). JHOC5 and OVCAR7 cells were generously provided by Drs. Evette Radisky (Mayo Clinic, Jacksonville, FL) and Viji Shridhar (Mayo Clinic, Rochester, MN), respectively. SKOV3 (#HTB-77), FT194 (#CRL-3445), and FT282 (#CRL-3449) cell lines were purchased from ATCC. MDAH, OVSAHO, COV362, SKOV3, OVCAR7, and JHOC5 cells were maintained in phenol red free RPMI medium (#11835050, Gibco) containing 10% (v/v) FBS (#900-308, Gemini Bioproducts) and 1% (v/v) antibiotic/antimycotic (#15240-062, Gibco). COV362-Olap^R^ cells were maintained as the parental line with the addition of 10 µM olaparib. PEO1, PEO4, and ABT^R^#2 cells were maintained in phenol red free DMEM/F12 medium (#41121800, Jango Bio) containing 10% (v/v) FBS, 1% (v/v) non-essential amino acids (#M7145, Corning), 10 µg/ml insulin (#IO516, Sigma-Aldrich), and 1% (v/v) Pen/Strep (#15070063, Invitrogen). FT194 and FT282 were maintained in phenol red free DMEM/F12 medium containing 10% (v/v) FBS, and 1% (v/v) Pen/Strep. All cells were cultured in a humidified 37°C incubator with 5% CO_2_. Cells were authenticated by IDEXX BioAnalytics (last tested March 2023) and tested for mycoplasma infection every 6 months using a Mycoplasma Detection Kit (#131001, SouthernBiotech).

### IncuCyte compatible cell line generation

OVSAHO, MDAH, PEO1, and ABT^R^#2 cells were infected at an MOI of 3-5 with the IncuCyte Nuclight Red (#4476 Sartorius) or IncuCyte Cell Cycle Green/Red Lentivirus Reagent (#4779, Sartorius), and pooled stably expressing cell lines were selected with and maintained in indicated medium further supplemented with 500 µg/L puromycin (#A11138, Gibco).

### Drug library screen

Indicated cell line models were plated at 2000 cells per well in 96 well plates (#3595, Corning). Twenty-four hours later, the 146 compounds contained within the Cayman epigenetics screening library, including DMSO as a vehicle control, were added at a 1 µM final concentration in duplicate. Following 48 hours of treatment, cell viability was determined using CellTiter-Glo (#G9243, Promega) following the manufacturer’s protocol and assayed using a GloMax luminometer (Promega). Luminescence values for each compound were averaged and normalized to the DMSO control wells to generate drug response values.

### Proliferation assays

Cells were plated at a density of 1000-2000 cell per well in 96 well plates in 100 μL of cell type specific medium and allowed to adhere overnight. For basal growth monitoring, indicated cells were placed in the IncuCyte S3 instrument (Sartorius) and allowed to proliferate for 5-7 days at which point growth was compared by confluence and/or red nuclei count at the same time point across each comparison. Dose–response curves were performed for indicated drugs using 9 rounds of 2-fold serial dilutions starting at 2.5 µM for lestaurtinib, 50 µM for ruxolitinib, olaparib, LY3214996, and CC-90001, or the dose range 50, 25, 10, 7.5, 5, 2.5, 1, 0.5 µM for cisplatin. Cells were maintained until vehicle treated wells neared confluence (between 5-10 days) at which point cells were fixed with 25% (v/v) glutaraldehyde (# G6257, Sigma-Aldrich) for 10 min, washed four times with water, stained with crystal violet (#C6158, Sigma-Aldrich), and washed again. Crystal violet-stained cells were solubilized with 100 µL of a solution containing 100 nM sodium citrate (#W302600, Sigma-Aldrich) and 50% ethanol and quantified using a plate reader at 550 nm excitation. IC50 values were calculated using a sigmoidal dose-response curve (log(inhibitor) vs normalized response -- Variable slope) in GraphPad Prism.

### Cell cycle profiling

Indicated lines expressing the cell cycle reporter were seeded in 96-well plates at a density of 2000-5000 cells per well and treated 24 hours later with DMSO or the indicated drugs. Immediately following treatment, cells were imaged every 2-4 hours in an IncuCyte S3 instrument (Sartorius) and the percent of cells in each cell cycle phase was calculated according to the manufacturer’s protocol.

### Apoptosis assays

Cells expressing Nuclight Red were seeded in 96-well plates at a density of 1000-2000 cells per well and treated 24 hours later with DMSO or the indicated drugs plus a 1:1000 dilution of Annexin V-Green (#4642, Sartorius) or a 1:2000 dilution of Caspase3/7-Green (#4440, Sartorius). Cells were imaged every 6 hours for 5-7 days in an IncuCyte S3 instrument and the ratio of apoptotic cells (green) to total cells (red) was determined.

### Colony formation assays

Cells were plated in triplicate at a density of 200 cells per well in 6- well plates and treated with the indicated agents for approximately 3 weeks. Colonies were stained with crystal violet as described above and the number of colonies was quantified.

### RNA-sequencing and analysis

Following treatment of cells with DMSO or lestaurtinib (500 nM) for 24 hours, total RNA was isolated using the Qiagen miRNeasy mini kit (#217004, Qiagen) following the manufacturer’s instructions. Library preparation and sequencing was performed by the Mayo Clinic Genome Analysis Core using an Illumina HiSeq 4000. Reads were aligned using STAR (version 2.7.3a) and featureCounts (version 2.8.2) was used to assign mapped reads. Any gene with an RPKM < 1 in all samples were considered to be lowly expressed and removed from further analysis. Data processing of the of PDX sequencing results was previously reported^14,15^. Comparison tests were performed using edgeR^16^ and significance was measured by |fold change| ≥ 1.5, P < 0.05 and FDR < 0.1. Ingenuity Pathway Analysis (IPA) software (Ingenuity Systems Inc.) was used to identify significantly altered biological pathways and upstream regulators from the differentially expressed genes (P < 0.05 and |z-score| > 2).

### PDX *ex-vivo* culture

Experiments were approved by the Institutional Animal Care and Use Committee (IACUC) at the Mayo Clinic. PDXs were developed at Mayo Clinic and maintained in female SCID mice as previously described^17^. Freshly harvested PDXs were dissociated and tumor cells, including tumor associated mouse cells, were plated in nano-culture plates (#NG-PLH9010, OrganoGenix) at a density of 40,000 cells per well and allowed to form spheroids for 3 days^18^. Spheroids were treated as indicated for 3 days and viability was measured using 3D CellTiter-Glo (#G9682, Promega) and a GloMax luminometer (Promega).

### Real-time qPCR

Cells were seeded at 50% confluence and treated as indicated, followed by total RNA isolation using TRIzol™ Reagent (#15596018, Qiagen). cDNA was generated from 1 µg of total RNA via an iScript™ cDNA Synthesis Kit (#1708891, Bio Rad), and subsequently used for real-time quantitative PCR (RT-qPCR) using PerfeCTa™ SYBR Green Fast Mix™ (#95071-012, Quanta Biosciences) and a Bio-Rad CFX Real-Time PCR detection system. Primers were purchased from Integrated DNA Technologies (IDT) and their sequences are listed in Supplemental Table 1.

### Western blots

Cells were seeded at approximately 70% confluence and treated as indicated for 30 mins. Cell lysates were generated using NETN buffer (150 mM NaCl, 1 mM EDTA, 20 mM pH 8.0 Tris, 0.5% NP-40) containing 1X cOmplete™ Protease Inhibitor Cocktail without EDTA (PI) (#11836170001, Roche) and 1X PhosSTOP™ phosphatase inhibitor (#049068370001, Roche). Twenty µg of protein was loaded into each lane of SDS-containing Criterion XT gels and western blotting was performed as previously described^19^. Antibody information is reported in Supplementary Table 2.

### siRNA knockdown

siRNAs designed to specifically target human STAT1 or STAT3 were purchased from Dharmacon (#L-003543-00-0010, # L-003544-00-0010). Cells were transfected with 5 nM ON-TARGETplus SMARTpool siRNAs using DharmaFECT 1 reagent (#T-2005-01, Dharmacon) according to the manufacturer’s protocol. Non-Targeting siRNA Pool 1 (#D-001206-13, Dharmacon) was used as the negative control.

### CRISPR knockout cell line generation

The STAT1 and STAT3 CRISPR guide RNAs (crRNAs, Supplemental Table 3) were designed using Synthego and the UCSC genome browser, where they were they were predicted as high-confidence guide RNAs.^20^ Guide RNAs, along with Alt-R CRISPR-Cas9-tracrRNA 5’ATTO550 (tracrRNA, #1075927) and Alt-R Cas9 protein (#1081058), were purchased from IDT. Cells were plated in 10 cm dishes at approximately 50% confluence. Twenty-four hours later, equimolar amounts of crRNA in 1 mL nuclease free water and tracrRNA in 1 mL duplex buffer were heated to 95⁰C for 5 minutes, allowed to cool to 21°C, and placed on ice for 30 minutes to form the sgRNA complex. The Lipofectamine™ CRISPRMAX™ Cas9 Transfection kit (#CMAXX00-001, Thermo-Fisher Scientific) was used to transfect Cas9 protein and the sgRNA complex into MDAH and OVSAHO cells following the manufacture’s protocol. The top 10% ATTO550 positive cells were collected by flow sorting and seeded in 10 cm dishes. Individual colonies were picked, expanded, and screened by western blotting. Clones with no detectable STAT1 or STAT3 protein expression were further screened at the genomic level via sanger sequencing.

### Drug Synergy

Cells were plated in 96 well plates at a density of 1000-2000 cells per well in triplicate and allowed to adhere overnight. Cells were then treated with monotherapy or combination therapy as indicated using 3-fold serial dilutions, with the concentration ranges determined by individual drug IC50s with the highest dose killing close to 100% of cells and the lowest dose killing close to 0% of cells. Following 5-10 days of treatment, cell viability was measured using CellTiter-Glo. Synergy was assessed using SynergyFinder^21^ and the Compusyn software^22^.

### Statistical analyses

All experiments were conducted in at least 3 biological replicates with 3-8 technical replicates per assay. Representative data sets are shown. Student’s t-test and two-way ANOVAs were used to determine significant differences between treatments and p-values <0.05 were considered statistically significant after adjusting for multiple comparisons (Tukey adjustment) when necessary. Graphical representations and statistical analyses were performed using GraphPad Prism.

### Data availability

The data generated in this study are publicly available in Gene Expression Omnibus (GEO) at GSE###### (we will provide the GSE number upon acceptance of our manuscript).

## Results

### Drug screen identifies lestaurtinib as an inhibitor of therapy-sensitive and -resistant ovarian cancer cells

To identify novel inhibitors of ovarian cancer cell viability, we performed a drug screen using 1 µM concentrations of a panel of small molecules targeting epigenetic regulators. To enhance generalizability of the results, we employed representative cell lines of the two most common ovarian cancer subtypes^23^, high-grade serous (OVSAHO) and endometrioid (MDAH) and their isogenic cisplatin-resistant clones^11^. Approximately 25% of the compounds were found to inhibit cell viability in at least 1 of the 4 cell lines (Figure 1A) and hits were prioritized as indicated in the schema (Figure 1B). Four compounds were identified that met these criteria: (S)-HDAC-42, SB939, CPI-203, and lestaurtinib. (S)-HDAC-42 and SB939 are both HDAC inhibitors and are part of a drug class that is widely studied in ovarian cancer^24^. CPI-203 is an epigenetic reader domain inhibitor that binds to the bromodomains of BRD4 and has previously been studied in the context of ovarian cancer^25^. In contrast, lestaurtinib (CEP-701, Teva), a broad spectrum tyrosine kinase inhibitor known to target FLT3, JAK2, and PRK1^26^, has never been studied in the context of ovarian cancer, thus we elected to further study this compound. To validate the antineoplastic activity of lestaurtinib, we conducted dose response cell viability assays in an expanded panel of ovarian cancer cell lines, including MDAH, MDAH cisplatin-resistant, OVSAHO, OVSAHO cisplatin resistant, SKOV3 (serous cystadenocarcinoma), OVCAR7 (high grade serous), JHOC5 (clear cell carcinoma), PEO1 (high grade serous), two isogenic therapy resistant PEO1-related clones (PEO4; cisplatin-resistant and ABT^R^#2; PARPi-resistant), COV362 (high grade serous), and its isogenic PARPi-resistant clone (COV362-Olap^R^). Across this panel, the lestaurtinib IC50 concentrations ranged from 10-410 nM (Figure 1C), which is far below the known C_max_ and C_trough_ values for lestaurtinib (7-12 µM and 1-12 µM, respectively) in patients receiving a standard dose of 80 mg twice daily^27,28^.

**Figure 1.**
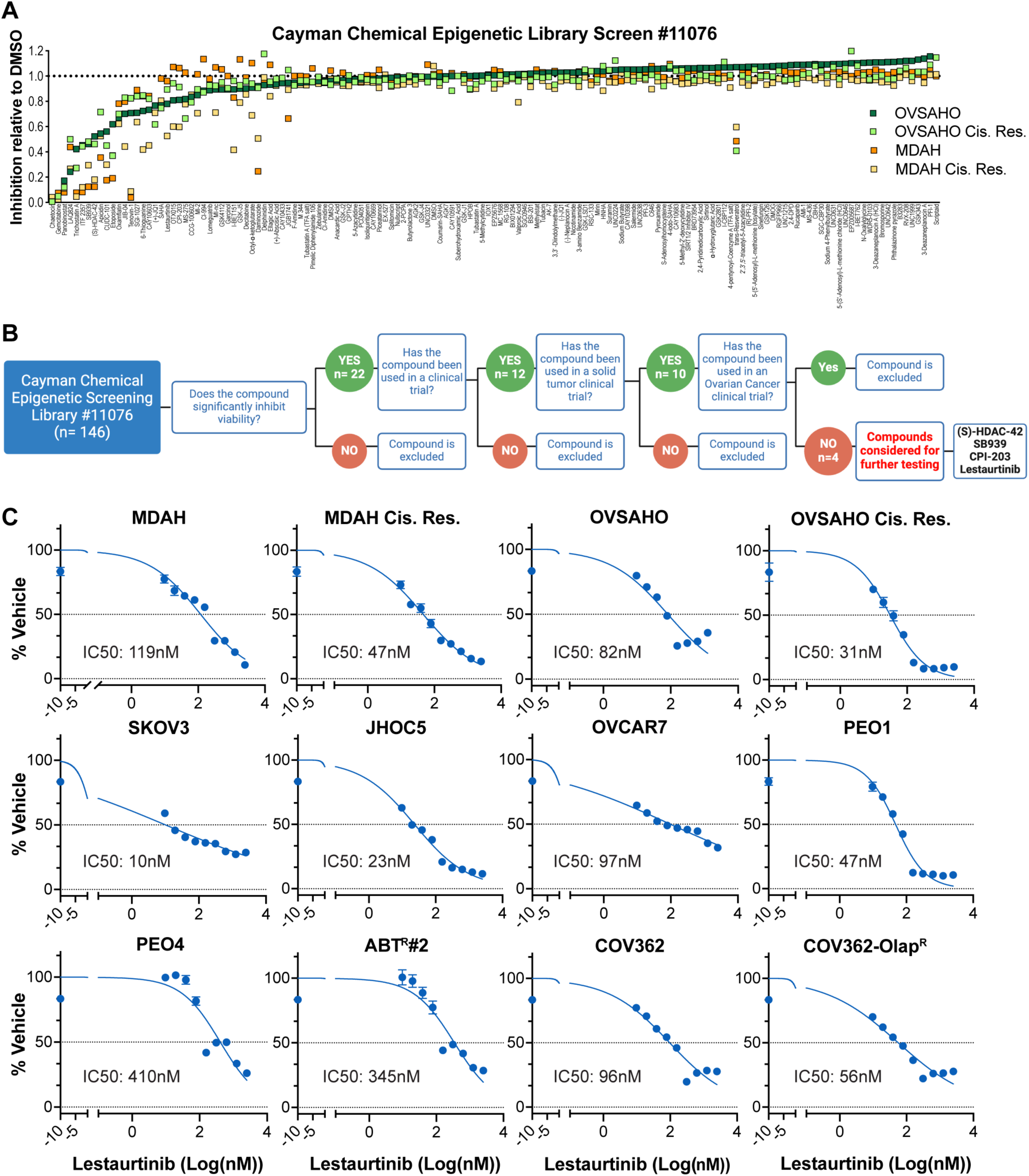
Identification of lestaurtinib as a novel inhibitor of sensitive and resistant ovarian cancer cells. **A)** Viability of indicated cell lines following 48 hours of treatment with 146 compounds (1 µM) contained in the Cayman Chemical epigenetic drug library (#11076) relative to DMSO vehicle. **B)** Drug screen hit prioritization schema. Figure created with BioRender.com. **C)** Dose response curves (2-fold dilution starting at 2.5 µM) and IC50 concentrations for lestaurtinib in a panel of sensitive and resistant ovarian cancer cell lines following 5-10 days of treatment. Abbreviations: Cis Res = cisplatin-resistant; Olap^R^ = olaparib-resistant.

### Lestaurtinib induces cell cycle arrest and apoptosis and inhibits colony formation

To determine whether lestaurtinib is cytostatic or cytotoxic, we conducted cell cycle profiling and assessed levels of apoptosis. OVSAHO, MDAH, PEO1, and ABT^R^#2 cell cycle reporter lines were treated with escalating doses of lestaurtinib. Following 24 hours of treatment, lestaurtinib induced a G1 cell cycle arrest with a corresponding G2/M-phase diminution in all 4 cell lines (Figure 2A; Supplemental Figure 2A). Longer treatment (2-5 days) with lestaurtinib induced substantial levels of apoptosis as measured by Caspase-3/7 and Annexin V reporter assays at these same concentrations (Figure 2B; Supplemental Figure 2B-C). Finally, lestaurtinib concentrations of ≥ 250 nM markedly inhibited colony formation in all cell models (Figure 2C; Supplemental Figure 2D). These results indicate that lestaurtinib rapidly induces cell cycle arrest followed by progressive induction of apoptosis, resulting in significant suppression of cell viability at clinically achievable plasma concentrations.

**Figure 2.**
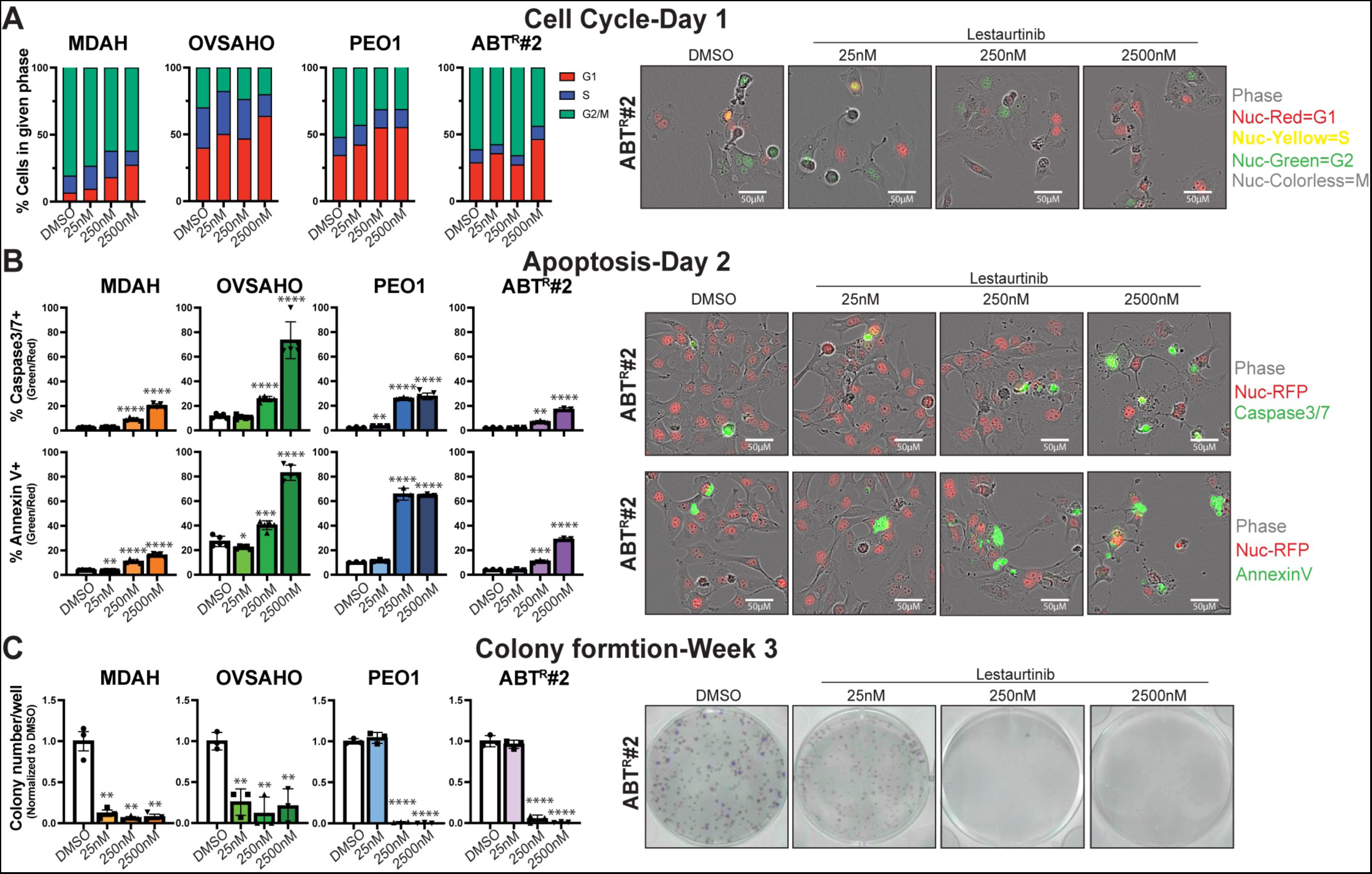
Lestaurtinib induces cell cycle arrest, apoptosis, and prevents colony formation. **A)** Indicated cell lines expressing a fluorescent cell cycle reporter were treated with multiple concentrations of lestaurtinib for 24 hours and cell cycle phase distribution was monitored. The percentage of cells in each phase of the cell cycle was quantified via the IncuCyte S3 system based on the presence of nuclear red (G1), nuclear green (G2/M) and nuclear yellow (S). Representative images of the ABT^R^#2 cell line are shown. **B)** Nuc Red expressing cells were incubated with Caspase 3/7-GFP or Annexin V-GFP reporters and subsequently treated with lestaurtinib for 2 days. Percent apoptotic cells were defined as # green positive cells relative to # of red positive cells. Representative images of the ABT^R^#2 cell line are shown. **C)** Colony formation was assessed in the indicated cell lines following 3 weeks of treatment with multiple concentrations of lestaurtinib and the number of colonies were quantified by crystal violet staining. Representative images of the ABT^R^#2 cell line are shown. Abbreviations: Nuc = nuclear. Graphs represent mean ± standard error. ANOVA p-values: *<0.0332,**<0.0021,***<0.0002,****<0.0001.

### Lestaurtinib inhibits JAK/STAT signaling

To elucidate the genome-wide effects of lestaurtinib on ovarian cancer cells, we conducted RNAseq in MDAH and OVSAHO cells treated with 500 nM lestaurtinib for 24 hours. Surprisingly, MDAH and OVSAHO cells exhibited striking differences in their response to lestaurtinib with 824 and 778 uniquely regulated genes, respectively (Figure 3A-D). Of the 161 commonly regulated genes between MDAH and OVSAHO cells, 61.5% were directionally concordant (Figure 3E). Interrogation of the differentially regulated genes by lestaurtinib in each cell line via IPA revealed significant alterations in many inflammatory pathways (IL-17 signaling, rheumatoid arthritis, neuroinflammation, acute phase response, S100 family signaling, cytokine production and signaling, TNFR1/2 signaling, inflammatory airway diseases, and iNOS signaling), which are all comprised of JAK/STAT signaling components (Figure 3F-G). Cell-cell communication pathways (tumor microenvironment, EMT by growth factors pathway, and neuregulin signaling), pro-cancer/proliferation pathways (cholestrerol biosynthesis, ceramide signaling, p53 signaling, glioma signaling), and HIF1α signaling were also impacted (Figure 3F-G). Upstream regulator analysis indicated that inhibition of STAT-mediated transcription likely contributed to the differential gene expression profiles following lestaurtinib treatment (Figure 3H-I). Taken together, these results indicate that lestaurtinib suppresses the JAK/STAT pathway in MDAH and OVSAHO cell lines.

**Figure 3.**
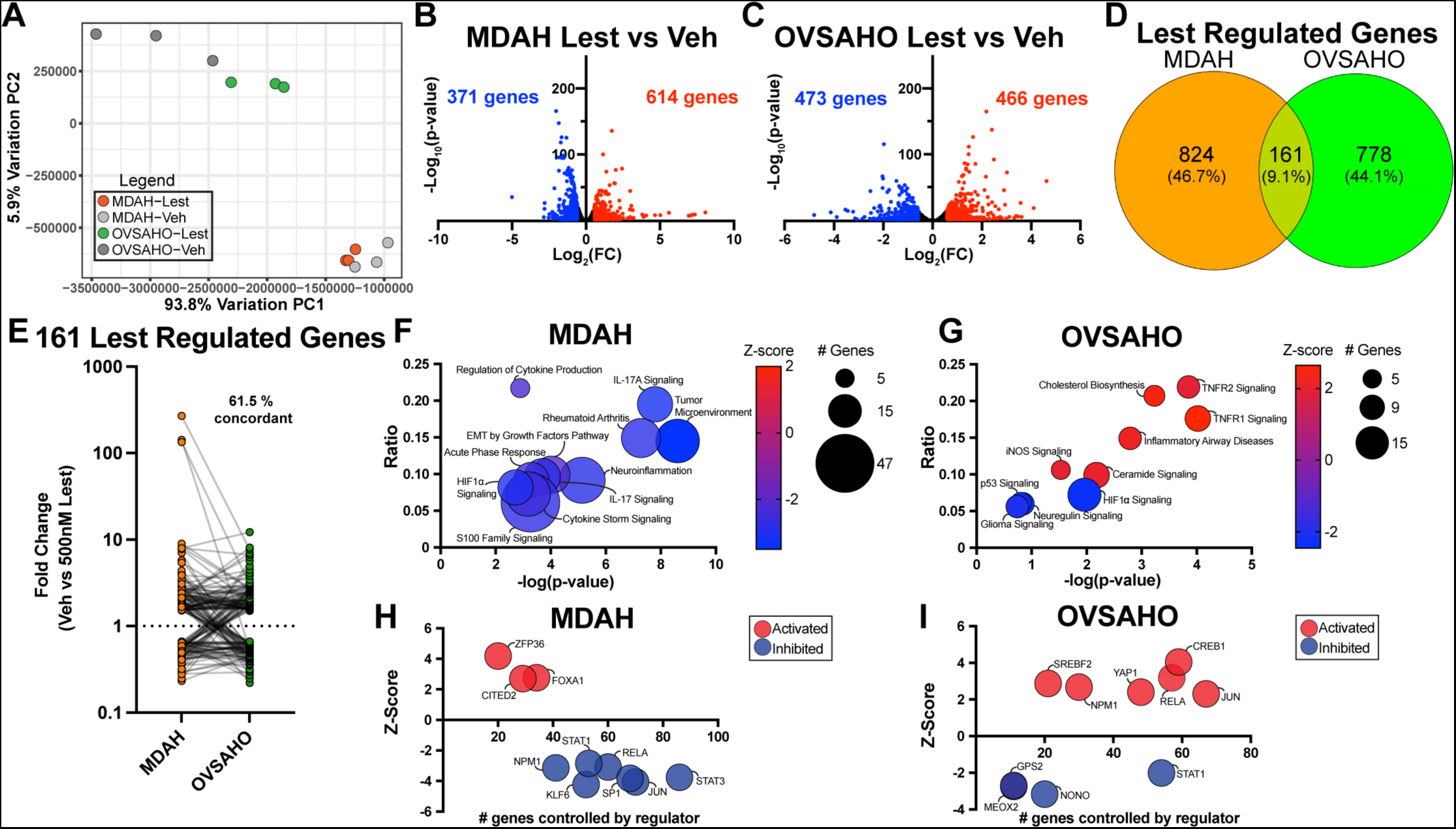
Lestaurtinib inhibits JAK/STAT signaling. RNAseq was conducted on MDAH and OVSAHO cells treated with 500 nM lestaurtinib or DMSO for 24 hours. **A)** PCA plot of resulting data from both cell lines. **B&C)** Volcano plots of differentially expressed genes in MDAH and OVSAHO cells where blue indicates down-regulated and red indicates up-regulated genes (|fold change| ≥1.5, p<0.05 and FDR<0.1). The number of significantly regulated genes are shown. **D)** Venn diagram depicting the overlap of lestaurtinib regulated genes in MDAH and OVSAHO cells. **E)** Plot depicting the concordance of lestaurtinib regulated genes that were commonly altered between MDAH and OVSAHO cell lines. **F&G)** Ingenuity Pathway Analysis and **H&I)** upstream regulator analysis of lestaurtinib regulated genes in MDAH and OVSAHO cells. Abbreviations: Lest = lestaurtinib, Veh = vehicle. Pathways and regulators with p<0.05 and |z-score|>2 are shown.

### Activity of lestaurtinib in ovarian cancer PDX models *ex vivo*

To further extend the potential clinical relevance of these observations in ovarian cancer, the inhibitory effects of lestaurtinib were assessed in a panel of ovarian cancer PDXs grown *ex vivo* as spheroid cultures (Figure 4A). Of the 8 models tested, 4 were responsive to lestaurtinib while another 4 were not (Figure 4B). Four of the PDX models represent PARPi-sensitive and -resistant pairs (Supplemental Table 4). PH039S and PH077S were derived from treatment naïve patients who responded to cisplatin treatment and have previously been confirmed to be sensitive to paclitaxel/carboplatin treatment both with and without niraparib^29^, as well as to carboplatin and niraparib as single agents^15^. PH039R and PH077R represent derivatives of these two models that were developed to be PARPi-resistant through 4 serial cycles of niraparib treatment in mice^15^. Of note, the PARPi-resistant PH039R derivative was exquisitely sensitive to lestaurtinib whereas the treatment naïve parental model was not (Figure 4B). In contrast, neither of the PH077 models responded to lestaurtinib (Figure 4B).

**Figure 4.**
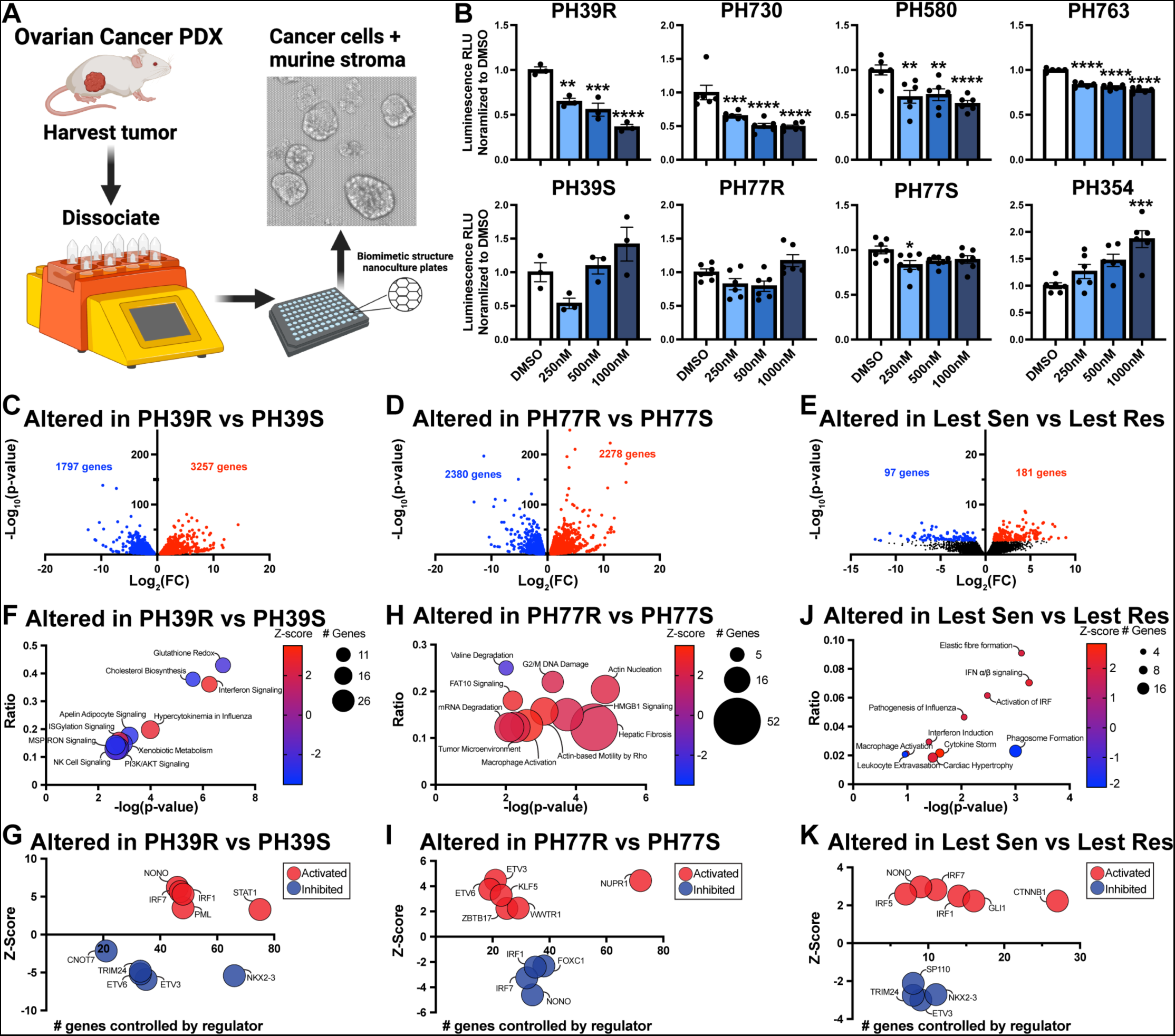
Efficacy of lestaurtinib in *ex vivo* PDX models. **A)** Depiction of ovarian cancer PDX processing for rapid evaluation of drug efficacy in *ex vivo* cultures. **B)** Viability of PDX models following 3 days of lestaurtinib treatment as assessed via 3D CellTiter-Glo. **C-E)** Volcano plots of differentially expressed genes in **C)** PH39R vs PH39S, **D)** PH77R vs PH77S, and **E)** lestaurtinib sensitive (PH39R, PH730, PH580) vs lestaurtinib resistant (PH39S, PH77S, PH77R, PH354) PDXs as determined by RNAseq. Blue represents significantly down-regulated genes and red indicates significantly upregulated-genes (|fold change| ≥1.5, p<0.05 and FDR<0.1). The number of significantly regulated genes are shown. **F-K**) Ingenuity Pathway Analysis and upstream regulator assessment of differentially expressed genes between the indicated PDX models. Pathways and up-stream regulators with p<0.05 and |z-score|>2 were considered as significant. Graphs depict mean ± standard error. ANOVA p-values: *<0.0332,**<0.0021,***<0.0002,****<0.0001.

Given these differences, we compared RNAseq data generated from these two tumor pairs, as well as 3 of the lestaurtinib-sensitive vs the 4 lestaurtinib-resistant models (Figure 4C-E). Differentially expressed genes in untreated PH039R vs PH039S were subjected to IPA which revealed that the top three upregulated pathways in PH039R were interferon signaling, hypercytokinemia in influenza, and ISGylation signaling, all of which rely on JAK/STAT signaling (Figure 4F). Additionally, STAT1 was identified as a top transcriptional regulator responsible for the induction of the more highly expressed genes in PH039R (Figure 4G). In contrast, none of the pathways or upstream effectors identified to be altered between untreated PH077R vs PH077S involved JAK/STAT signaling (Figure H-I). Further, when comparing lestaurtinib-sensitive vs - resistant PDXs, we discovered substantial upregulation of JAK/STAT related pathways including interferon induction, IRF activation, IFNα/β signaling, and cytokine storm signaling, with IRF1, 5, and 7 as identified upstream regulators (Figure 4J-K) in models that responded to lestaurtinib vs those that did not. These findings, in addition to those reported in Figure 3, strongly suggest that the antineoplastic activity of lestaurtinib may only be observed in cells/tumors with high JAK/STAT signaling and inflammatory pathway activity.

### JAK/STAT pathway activation in ovarian cancer cell lines

Considering the observation that lestaurtinib suppresses genes involved in JAK/STAT signaling and elicits potent antineoplastic activity in PDX models with highly active JAK/STAT signaling, we characterized key components of this pathway across a panel of ovarian cancer cell lines for which isogenic treatment-resistant derivatives were available. At the RNA level, JAK1 was the most abundant of the kinases, while STAT1 and STAT3 were the predominant STATs detected (Figure 5A). At the protein level, JAK1 and JAK2 were robustly expressed, while JAK3 was only detected in one of the ovarian cancer cell lines (Figure 5B). JAK2 exhibited similarly high expression in the two immortalized fallopian tube epithelial cell lines, while JAK1 was much less abundant relative to cancer cells (Figure 5B). Total and phosphorylated forms of STAT1, STAT3 and STAT5 protein were highly variable, but were universally detected, apart from STAT5 in OVSAHO and the PEO resistant models (Figure 5B).

**Figure 5.**
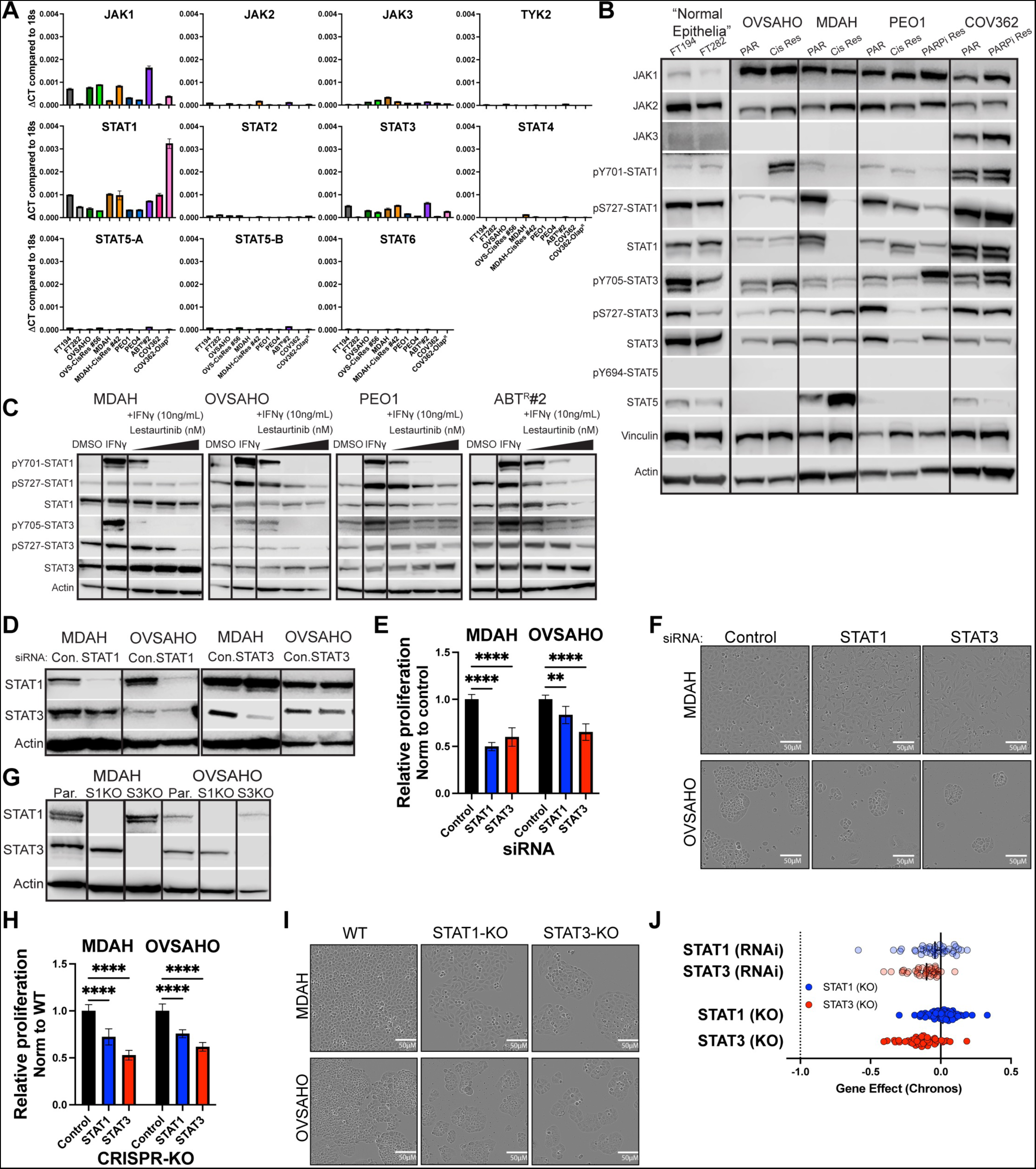
STAT1 and STAT3 are constitutively active and growth promoting in ovarian cancer cells. **A)** Relative mRNA expression levels of primary JAK/STAT family members in normal fallopian tube epithelial cells (FT194, FT282) and ovarian cancer cell lines under normal growth conditions. **B)** Western blots depicting the total protein and phosphorylation levels of primary JAK/STAT signaling components in indicated cell lines under normal growth conditions. Actin and vinculin served as loading controls. **C)** Western blots depicting the levels of STAT1 and STAT3 phosphorylation in response to 10 ng/mL IFNγ and increasing concentrations of lestaurtinib for 30 minutes. Actin served as the loading control. **D)** Western blots depicting total STAT1 and STAT3 protein following 72-hour treatment with scrambled control, STAT1 or STAT3 siRNA. Actin served as the loading control. **E)** Relative cell proliferation rates in response to siRNA-mediated STAT1 or STAT3 knockdown 5 days after transfection. **F)** Representative brightfield images depicting cell density 5 days post-transfection. **G)** Western blots depicting total STAT1 and STAT3 protein levels in MDAH and OVSAHO parental, STAT1 (S1) and STAT3 (S3) CRISPR knockout (KO) cells. Actin served as the loading control. **H)** Relative proliferation rates of control, STAT1 and STAT3 KO cells and **I)** representative brightfield images depicting cell density 5 days after plating. **J)** STAT1 and STAT3 essentiality in a panel of 59 ovarian cancer cell lines assessed via RNAi or CRISPR KO screens from the DepMap database. Gene Effect <0 indicates dependency, <-1 indicates essentiality. Abbreviations: Par = parental, Cis Res = cisplatin resistant, PARPi-Res = PARPi-resistant, Con = control, WT = wildtype, KO = knockout. Graphs depict mean ± standard error. ANOVA p-values: *<0.0332,**<0.0021,***<0.0002,****<0.0001.

In further studies, we assessed the effects of cytokines and lestaurtinib on STAT1 and STAT3 activation in ovarian cancer cells. IFNγ robustly induced pY701-STAT1, pS727-STAT1, and pY705-STAT3 protein levels (Figure 5C). Little to no impact on pS727-STAT3 was observed, but this residue was found to be constitutively activated across all models (Figure 5C). Lestaurtinib treatment inhibited IFNγ-mediated phosphorylation of STAT1 and STAT3 in a dose dependent manner and largely suppressed pS727 phosphorylation below basal levels for both STAT1 and STAT3 (Figure 5C). These results indicated that JAK1 and JAK2 are the primary upstream effectors of the JAK/STAT pathway in ovarian cancer cells and that lestaurtinib effectively inhibits phosphorylation of two key residues on STAT1 and STAT3 that are required for their activity.

### Contribution of STAT1 and STAT3 to ovarian cancer cell viability and proliferation

To directly assess the roles of STAT1 and STAT3 in ovarian cancer cell viability and proliferation, siRNAs were used to selectively knockdown STAT1 and STAT3 in MDAH and OVSAHO cells (Figure 5D). Following STAT1 or STAT3 knockdown, MDAH and OVSAHO cells exhibited significant growth inhibition (Figure 5E-F). Given that siRNA-mediated knockdown of STAT1 and STAT3 were incomplete, we used CRISPR-Cas9 to interrupt the genes encoding these transcription factors. Successful knockouts were confirmed by western blotting and DNA sanger sequencing (Figure 5G & Supplemental Figure 3). Both STAT1-KO and STAT3-KO isogenic cells grew significantly slower than their respective parental controls (Figure 5H-I). These findings are in agreement with DepMap Portal^30^ data of 59 ovarian cancer cell lines indicating that knockdown or deletion of STAT1 and STAT3 are largely growth inhibitory (gene effect scores = -0.5-0), but not lethal (gene effect scores ≤ -1, Figure 5J). These genetic loss of function studies further implicate STAT1 and STAT3 as drivers of ovarian cancer cell proliferation.

### Lestaurtinib synergizes with standard-of-care agents

Because complete ablation of STAT1 or STAT3 was not sufficient to kill ovarian cancer cells, we next sought to determine whether inhibition of JAK/STAT signaling would enhance the cytotoxicity of standard-of-care therapies. We designed synergy studies based on the individual drug IC50s (Figure 1 and Supplemental Figure 4), altered the ratios between the two inhibitors, and used both the Loewe synergy score (synergy >10)^21^ and the Chou-Talalay combination index (CI, synergy <1 and highly synergistic <0.3)^22^ to assess synergistic effects. Lestaurtinib was found to synergize with cisplatin in all cell lines tested via the Loewe synergy score, though PEO1 cells (already platinum sensitive due to a BRCA2 mutation) exhibited a lower degree of synergy (Figure 6A), while the CI method identified synergy across all dose ratios tested in all cell lines, with highly synergistic combinations for MDAH and ABT^R^#2 cells (Figure 6B). Lestaurtinib also synergized with the PARPi olaparib across all cell lines according to the Loewe synergy score (Figure 6C). The CI method indicated that there were multiple dose ratios of lestaurtinib and olaparib that were strongly synergistic in MDAH, OVSAHO, and PEO1 cells (Figure 6D). Importantly, lestaurtinib was found to synergize with olaparib across multiple dose ratios in the ABT^R^#2 cells, a PARPi-resistant cell line, thus revealing its potential to resensitize cells to PARPi therapy which is a major unmet clinical need (Figure 6D). These data support the potential utility of lestaurtinib, not only as a monotherapy or salvage strategy, but also as a component of effective combination therapy, even in the setting of advanced and resistant forms of disease.

**Figure 6.**
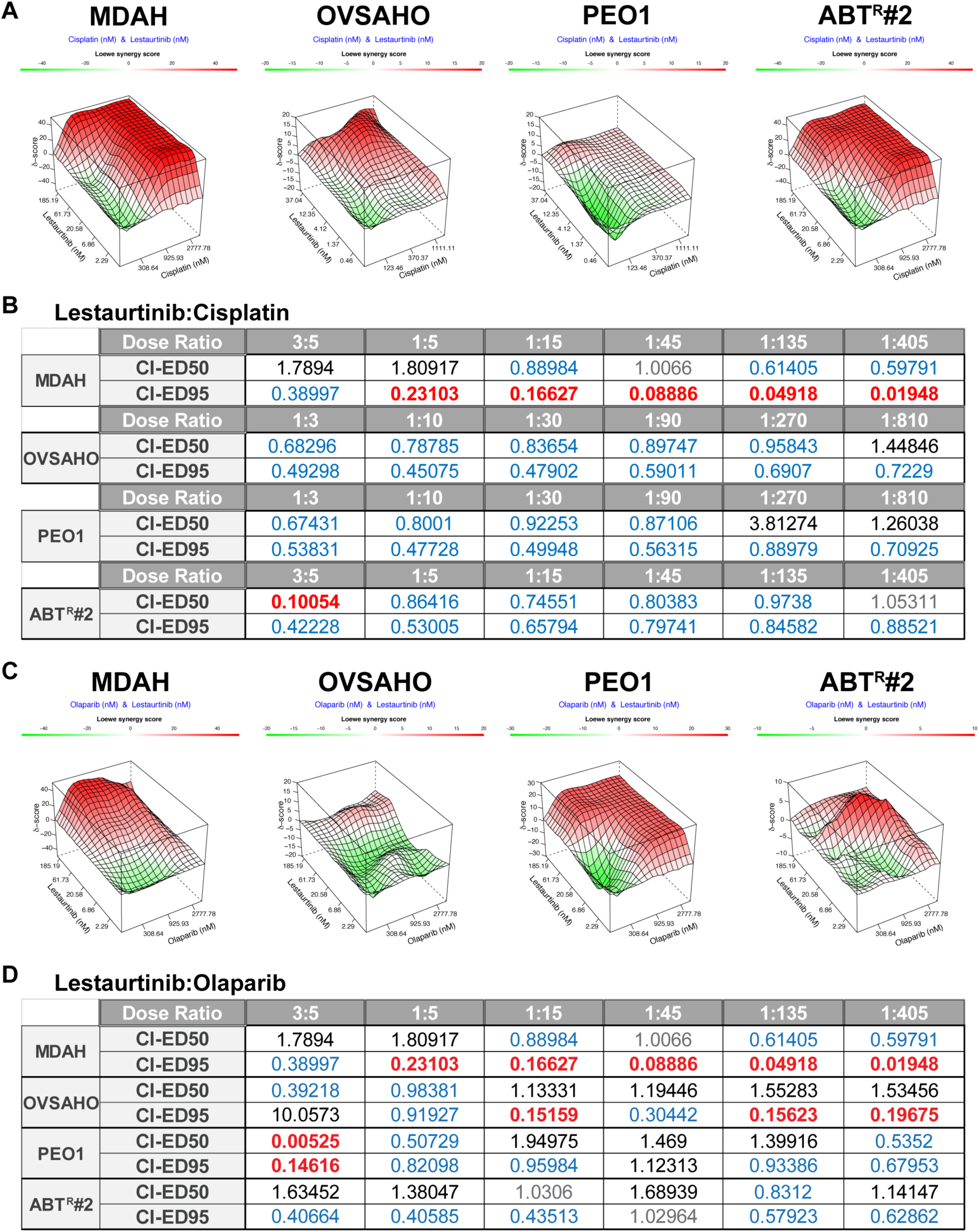
Lestaurtinib synergizes with cisplatin and olaparib. Indicated cells were treated with cisplatin (25000 nM or 10000 nM, 3-fold dilution) or olaparib (25000 nM, 3-fold dilution), alone and in combination with lestaurtinib (5000 nM or 1000 nM, 3-fold dilution), for 5-7 days. Relative cell viability was measured with crystal violet staining. **A&C)** Synergy plots depicting Loewe scores generated using SynergyFinder3.0. **B&D)** Combination index (CI) scores were calculated using the Chou-Talalay method and compusyn software. Antagonism = >1 (black), additive = 1 (grey), synergy = < 1 (blue), and highly synergistic = < 0.3 (red).

### Comparison of lestaurtinib and ruxolitinib in ovarian cancer cell lines

At present, the most well characterized JAK/STAT inhibitor is ruxolitinib^31^. To determine whether ruxolitinib would induce similar effects to that of lestaurtinib, we performed cell viability assays and derived IC50 values in our panel of ovarian cancer cell lines. Ruxolitinib IC50 values were near or above 20 µM (Figure 7A), a concentration far above the plasma C_max_ of 1 µM and a C_trough_ of 100 nM^32^ achieved in humans receiving the standard dose of 25 mg twice daily^31^. Similarly, ruxolitinib did not significantly inhibit any of the 8 PDX models treated *ex vivo* (Figure 7B), including those models with high levels of JAK/STAT pathway activity (Figure 4). Ruxolitinib did not induce cell cycle arrest after 24 hours of treatment nor did it impact apoptosis after 5 days of treatment (Supplemental Figure 5A-B). However, ruxolitinib did partially inhibit colony formation following 3 weeks of treatment (Supplemental Figure 5C).

**Figure 7.**
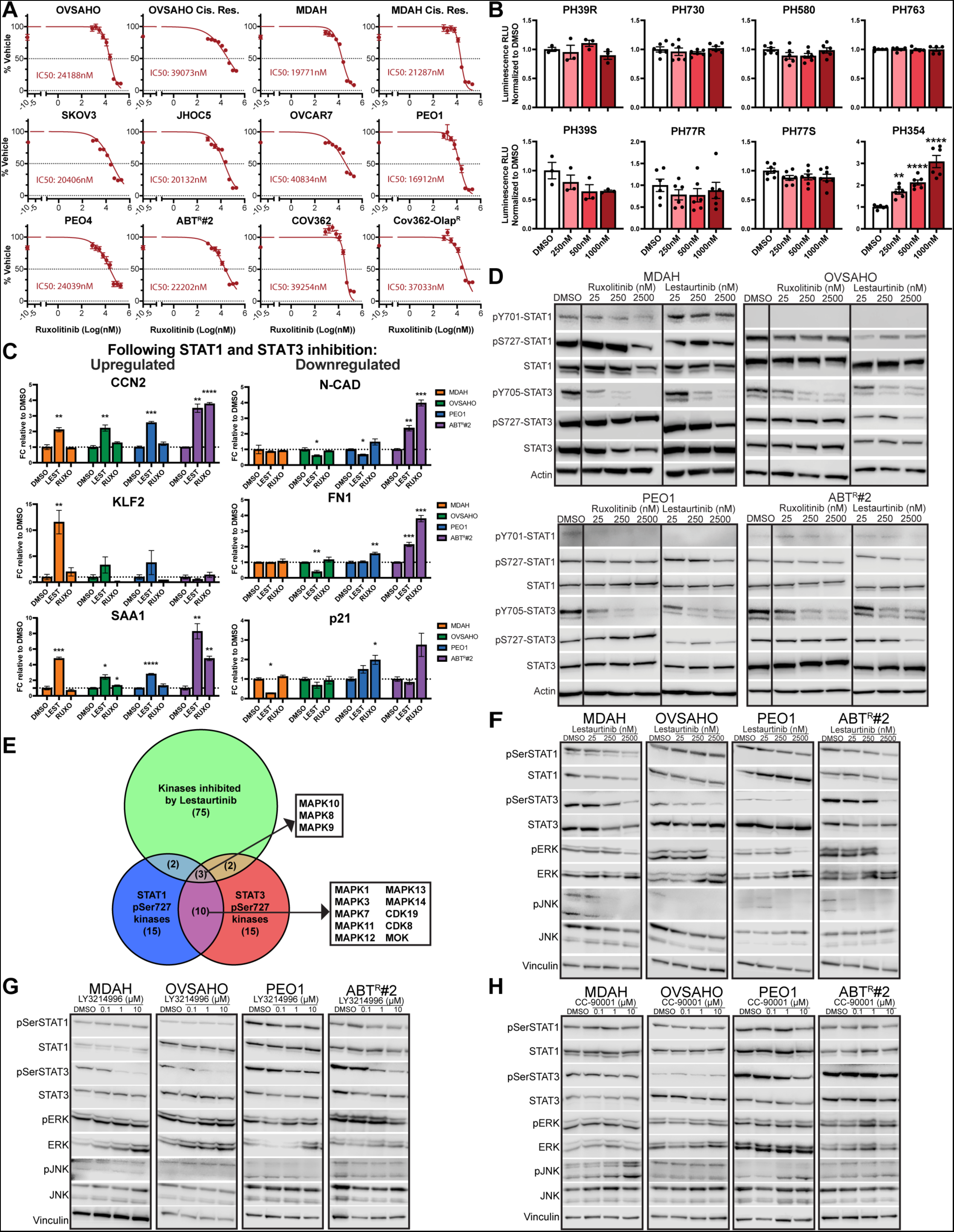
Mechanistic basis for superiority of lestaurtinib vs ruxolitinib. **A)** Dose response curves (2-fold dilution starting at 200 µM) and IC50 concentrations for ruxolitinib in a panel of sensitive and resistant ovarian cancer cell lines following 5-10 days of treatment. **B)** Viability of PDX models following 3 days of ruxolitinib treatment *ex vivo* as assessed via 3D CellTiter-Glo. **C)** Expression of indicated STAT1 and STAT3 regulated genes following 24 hours of treatment with DMSO, 250 nM lestaurtinib or 250 nM ruxolitinib. **D)** Western blots depicting the levels of STAT1 and STAT3 phosphorylation following treatment with DMSO or indicated concentrations of lestaurtinib and ruxolitinib for 30 minutes. Actin served as the loading control. **E)** Venn diagram of kinases known to be inhibited by lestaurtinib and predicted to phosphorylate S727 of STAT1 and/or STAT3. Western blots depicting total and phosphorylated levels of STAT1, STAT3, ERK and JNK following 30 minutes of **F)** lestaurtinib, **G)** LY3214996 or **H)** CC-90001 treatment. Vinculin served as the loading control. Abbreviations: Cis Res = cisplatin resistant, Lest = lestaurtinib, Ruxo = ruxolitinib. Graphs depict mean ± standard error. ANOVA p-values: *<0.0332,**<0.0021,***<0.0002,****<0.0001.

To examine the basis for the disparate effects of ruxolitinib and lestaurtinib, we assessed the ability of these two drugs to alter the expression of randomly selected genes identified to be downstream of STAT1 and STAT3 in our RNAseq studies (Figure 3). Following lestaurtinib treatment for 24 hours, CCN2 (4/4 cell lines), KLF2 (3/4 cell lines), and SAA1 (4/4 cell lines) gene expression was increased as expected, while ruxolitinib did not induce these genes with the exception of CCN2 and SAA1 in ABT^R^#2 cells (Figure 7C). For down regulated genes, lestaurtinib treatment suppressed N-CAD (3/4), FN1 (1/4), and p21 (3/4) while ruxolitinib elicited no inhibitory effects and instead actually increased expression of these genes in the PEO1 and ABT^R^#2 cells (Figure 7C). Further, we compared the ability of lestaurtinib and ruxolitinib to inhibit phosphorylation of the Y701/705 and S727 residues of STAT1 and STAT3. As shown in Figure 5C, lestaurtinib suppressed phosphorylation of these residues in a dose dependent manner. In contrast, ruxolitinib potently inhibited Y701/705 phosphorylation, but had little to no effect on S727 (Figure 7D). These results unveil a substantial difference between lestaurtinib and ruxolitinib regarding their effects on the phosphorylation profiles, and thus the activities, of STAT1 and STAT3 in ovarian cancer cells.

### Basis for lestaurtinib-mediated inhibition of STAT1 and STAT3 S727 phosphorylation

Given the ability of lestaurtinib to inhibit S727 phosphorylation of STAT1 and STAT3, we sought to identify the mechanistic basis for this inhibition. Kinases found to bind lestaurtinib with a K_D_ of ≤ 500 nM (82 total) were curated from a Kinobead screen^33^. We then interrogated the PhosphoSitePlus database to identify kinases predicted or known to phosphorylate the S727 residue on STAT1 and STAT3^34^ and pulled the top 30 kinases from each of these lists. When comparing the overlap of these lists (Supplementary Table 5), we found three kinases (MAPK8, MAPK9, and MAPK10 encoding JNK1-3) that were identified in all 3 datasets (Figure 7E). Ten additional kinases, predominated by the ERK family (MAPK1, MAPK3, MAPK7 and MAPK12), were commonly predicted to phosphorylate S727, but have not yet been shown to bind lestaurtinib (Figure 7E). Based on these findings, we assessed the effects of lestaurtinib on JNK and ERK via western blotting in a panel of ovarian cancer cells. Lestaurtinib not only inhibited S727 phosphorylation as previously shown (Figure 5C) but also suppressed phosphorylation of ERK1/2 and JNK1/2 in a dose-dependent manner (Figure 7F). Given these findings, we next assessed the effects of selective ERK (LY3214996) and JNK (CC-90001) inhibitors on ovarian cancer cell line growth and STAT1 and STAT3 S727 phosphorylation. The IC50 values for LY3214996 and CC-90001 ranged from 2-6 µM and 2-20 µM, respectively (Supplemental Figure 6). As shown in Figure 7G-H, both inhibitors significantly inhibited STAT3 S727 phosphorylation in a dose dependent manner, with some inhibitory effects on STAT1 in these ovarian cancer cell lines. Of note, LY3214996 and CC-90001 have both been shown to increase and decrease the phosphorylation levels of their respective targets depending on the cellular context^35–37^. In our ovarian cancer models, neither inhibitor had any impact on the phosphorylation of JNK or ERK (Figure 7G-H). Given the selective inhibition of LY3214996 and CC-90001 on S727 phosphorylation, we determined whether their combination with ruxolitinib, a potent inhibitor of Y701/705 phosphorylation, would recapitulate the effects of lestaurtinib. Interestingly, multiple dose ratios of ruxolitinib in combination with LY3214996 or CC90001 were found to be synergistic in all cell lines tested (Figure 8A-D). These findings further support the importance of blocking both Y701/705 and S727 phosphorylation on STAT1 and STAT3 for robust inhibition of ovarian cancer cell viability, an effect achieved by lestaurtinib monotherapy, but not by any of the other monotherapies examined here.

**Figure 8.**
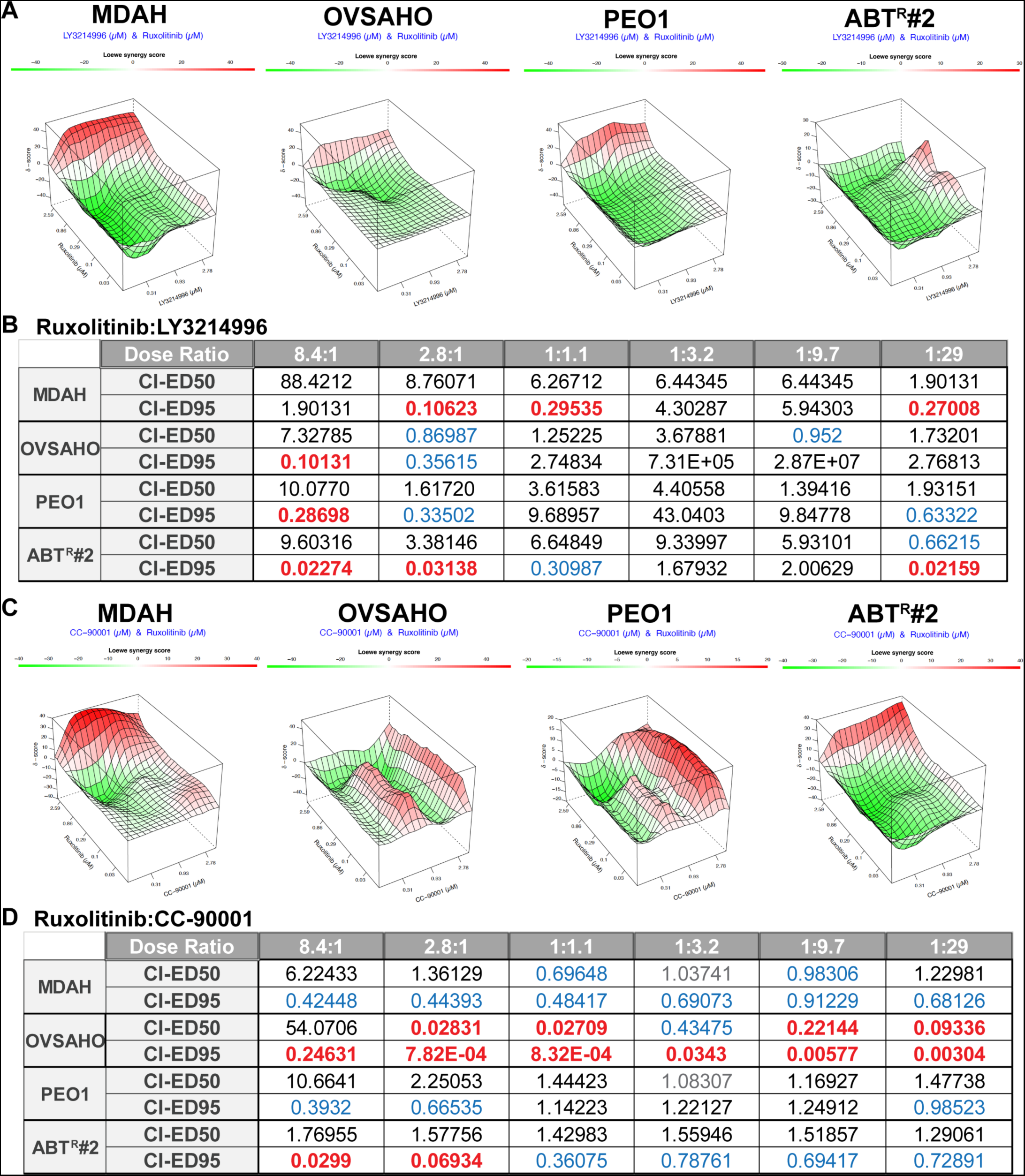
Ruxolitinib synergizes with LY3214996 and CC-90001. Indicated cells were treated with LY3214996 (70 µM, 3-fold dilution) or CC-90001 (70 µM, 3-fold dilution), alone and in combination with ruxolitinib (25 µM, 3-fold dilution) for 5-7 days. Relative cell viability was measured with crystal violet staining. **A&C)** Synergy plots depicting Loewe scores were generated using SynergyFinder3.0. **B&D)** Combination index (CI) scores were calculated using the Chou-Talalay method and compusyn software. Antagonism = >1 (black), additive = 1 (grey), synergy = < 1 (blue), and highly synergistic = < 0.3 (red).

## Discussion

In the present study, we identified lestaurtinib as a potent inhibitor of ovarian cancer cells with low nM IC50 values that are substantially less than clinically achievable plasma concentrations. We demonstrate that lestaurtinib uniquely and robustly inhibits JAK/STAT signaling through blockade of both Y701/705 and S727 phosphorylation of STAT1 and STAT3. Lestaurtinib treatment of PDX models *ex vivo* confirmed antineoplastic activity in 4 of 8 models tested. Intriguingly, high JAK/STAT pathway activity was predictive of lestaurtinib efficacy across these models. Moreover, lestaurtinib synergized with both cisplatin and olaparib, even in PARPi-resistant cells. Conversely, the most well-known JAK/STAT inhibitor, ruxolitinib, failed to suppress S727 phosphorylation of STAT1 and STAT3 and displayed no efficacy as a monotherapy in any cell line or PDX model tested. Intriguingly, when ruxolitinib was combined with either a JNK or ERK inhibitor, which were shown to inhibit S727 phosphorylation, synergistic antineoplastic activity was observed. Collectively, these observations have defined novel mechanisms of action by which lestaurtinib functions and highlight the potential of blocking multiple kinases upstream of STAT1 and STAT3 for the treatment of ovarian cancer.

The STAT family of transcription factors consists of seven members that share similar domain structures, including the transactivating domain (TAD) where Y701/705 and S727 are located. The TAD domain is essential for STAT transcriptional activity as it provides binding sties for essential co-regulators and stabilization at target gene promoters^38^. Phosphorylation of the tyrosine residue of both STAT1 and STAT3 is primarily mediated by JAKs, which are complexed with various cytokine receptors in response to cytokine mediated activation^39^. However, these tyrosine residues can also be directly phosphorylated by other receptor tyrosine kinases such as EGFR, PDGFR, and CSF1R, among others^40^. Upon phosphorylation of Y701/705, STAT proteins homo- or hetero-dimerize and translocate into the nucleus where they regulate gene expression^40^. The consequences of S727 phosphorylation are less well documented but have been reported to play a role in recruitment of transcriptional machinery^41,42^. Mutation of S727 to alanine, which prevents phosphorylation, has no impact on Y701/705 phosphorylation or STAT DNA binding *in vitro* or *in vivo*, but is necessary for maximal transcriptional output^43^. Further, blockade of STAT3 S727 phosphorylation has been shown to prevent DNA association of STAT3 homodimers, but not STAT1/STAT3 heterodimers or STAT1 homodimers^44^. These observations provide a potential explanation for the importance of inhibiting S727 phosphorylation in the present study.

JAK/STAT signaling is reported to be oncogenic in many forms of cancer, including ovarian cancer^45,46^, and is constitutively active in a proportion of chemotherapy resistant ovarian cancer cell lines^47^ and tumors^48^. STAT3 is widely regarded to promote progression of cancer, while the role of STAT1 is less clear^49,50^. A recent study utilizing single cell RNA-sequencing of ovarian tumors and PDX models reported that JAK1, STAT1 and STAT3 were expressed in 80-90% of all tumor cells and were among the most highly abundant transcripts within those cells^51^. Further, JAK/STAT activating cytokines were among the most highly expressed genes in cells of the tumor microenvironment^51^ and have previously been linked to chemotherapy resistance and decreased progression free survival^52,53^. Considering the abundance of JAK/STAT signaling pathway components in ovarian cancer patient tumors, the ability of lestaurtinib as a monotherapy to robustly inhibit this pathway highlights its potential clinical utility.

Ruxolitinib, which is FDA approved for the treatment of myelofibrosis, polycythemia and graft-versus-host disease^31^, is also the only JAK/STAT inhibitor that has been evaluated in a clinical trial for ovarian cancer (NCT02713386). This recently completed phase I/II study identified the maximum tolerated dose of ruxolitinib in the presence of standard-of-care paclitaxel and carboplatin, and subsequently evaluated the efficacy of this combination on PFS relative to paclitaxel/carboplatin alone^54^. Ruxolitinib was found to be well tolerated with acceptable toxicity and offered a modest improvement in PFS of 3 months (14.6 months vs 11.6 months) with a hazard ratio of 0.702 (p=0.059) without any improvement in overall survival (HR=0.785; p=0.70) in the neoadjuvant setting for patients with newly diagnosed stage III-IV ovarian/fallopian tube/primary peritoneal carcinoma^54^. In the present report, we demonstrate that ruxolitinib does not inhibit the viability or proliferative capacity of ovarian cancer cells and PDX models, even at concentrations much higher than can be clinically achieved. Nevertheless, the recent ruxolitinib trial has demonstrated the feasibility of implementing early-phase randomized studies for front line treatment of ovarian cancer and suggests that similar trials with more potent JAK/STAT inhibitors should be considered.

It is important to note that multiple classes of drugs have been designed to disrupt JAK/STAT signaling by targeting different nodes of this complex pathway. These include antibodies to block specific cytokines or their cognate receptors, such as interleukins and TNF, which are used to treated rheumatoid arthritis, inflammatory bowel disease and psoriasis^55^; small molecules similar to ruxolitinib, aimed at broadly or selectively inhibiting specific JAK proteins^56^; peptidomimetics targeting the SH2 domains of specific STATs in order to prevent dimerization and/or nuclear translocation^55^; computationally designed non-peptide based small molecules that can selectively bind to STAT proteins and disrupt their transcriptional output^55^; and nucleotide-based molecules, such as decoy oligonucleotides, antisense oligonucleotides, and siRNAs that disrupt STAT protein-DNA interaction or cause STAT protein downregulation^55^. The present study not only adds lestaurtinib, an agent with a well understood safety profile, to the list of JAK/STAT pathway inhibitors, but also demonstrates the ability of lestaurtinib to inhibit S727 as well as Y701/705 phosphorylation, which appears to contribute to its antineoplastic activity in ovarian cancer models.

While the present study has demonstrated important roles of STAT1 and STAT3 in ovarian cancer progression and has found that blockade of both Y701/705 and S727 is necessary to achieve potent antineoplastic effects, additional studies are needed to evaluate the exact mechanism by which these transcription factors drive cancer progression. First, it is not yet known how differential phosphorylation of various STAT1 and STAT3 sites contributes to their oncogenic activity and future studies of phosphomutant and phosphomimetic versions of STAT1 and STAT3 are needed to address this question. Second, tumor associated macrophages and fibroblasts in the ovarian tumor microenvironment not only produce JAK/STAT activating cytokines, but also exhibit high expression of JAK1, STAT1, and STAT3^51^. Future studies are needed to delineate the effects of lestaurtinib and other JAK/STAT inhibitors on STAT phosphorylation in these cell types, the resulting consequences on the tumor microenvironment, and ultimately the combined impact on tumor biology and clinically relevant outcomes. Such understanding will allow for the rational design and development of highly specific molecules that precisely disrupt STAT-mediated oncogenic cascades, while potentially sparing essential functions in anti-tumor immune cells and normal tissues. Finally, our RNAseq data revealed that lestaurtinib also impacts other pathways, which remain to be further explored to determine their contributions to lestaurtinib’s antineoplastic activity.

In summary, we have shown that JAK1, STAT1, and STAT3 are among the most abundantly expressed components of the JAK/STAT pathway in ovarian cancer cells; that constitutive activation of STAT1 and STAT3 frequently occur in ovarian cancer cells and PDX models; that lestaurtinib is a potent inhibitor of JAK/STAT signaling with the ability to block multiple phosphorylation events that are essential for STAT transcriptional activity; and that blockade of these phosphorylation events leads to cell cycle arrest followed by high levels of apoptosis that result in lestaurtinib-mediated killing of ovarian cancer cells, especially in settings where JAK/STAT signaling is elevated or constitutively active. These observations provide support for lestaurtinib, and other inhibitors that converge on this pathway, to undergo further preclinical testing and potentially future clinical study for the treatment of highly aggressive and refractory forms of ovarian cancer.

---

**Supplemental Figure 1.**
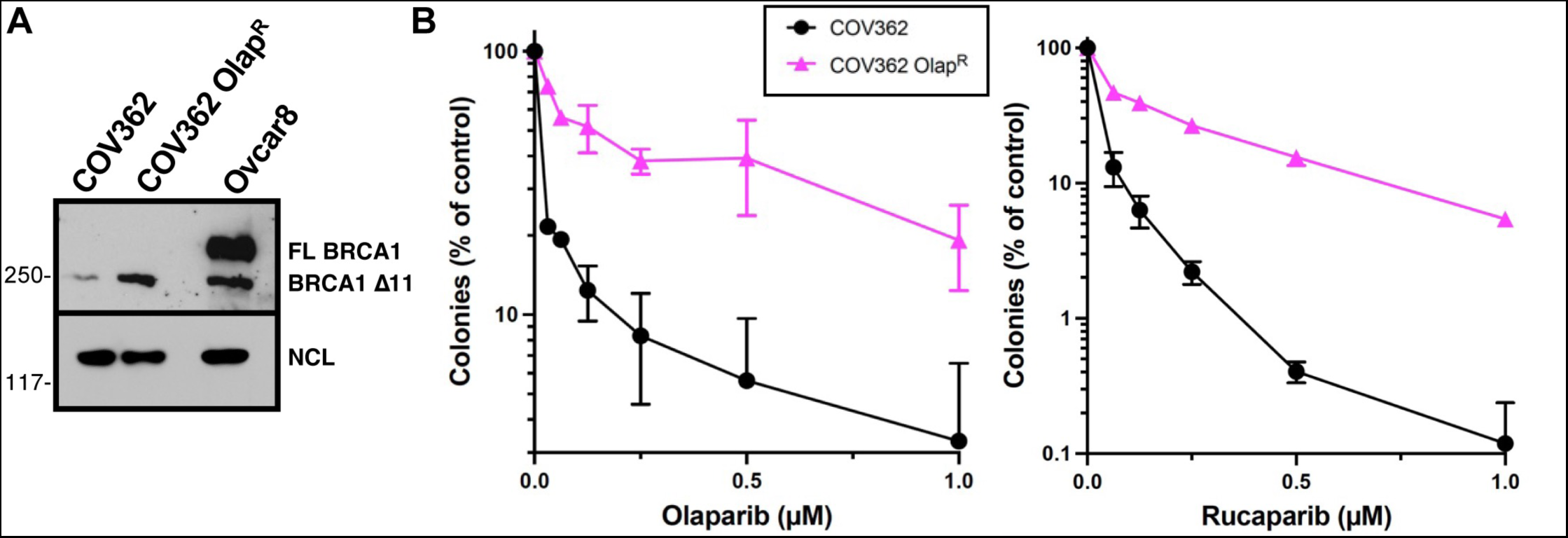
BRCA1 blotting and drug sensitivity of COV362-Olap^R^ cells. Parental COV362 cells were selected in increasing olaparib concentrations up to 10 µM. **A)** Western blot of nuclear proteins were probed with anti-BRCA1. Ovcar8 cells, which express wildtype BRCA1 were included as a positive control. Migration of full length (FL) BRCA1 and the Δ11 splice variant are shown. NCL served as a loading control. **B)** Parental and olaparib-resistant COV362 cells were allowed to form colonies for 14 days in the presence of the indicated concentrations of olaparib (left) or rucaparib (right).

**Supplemental Figure 2.**
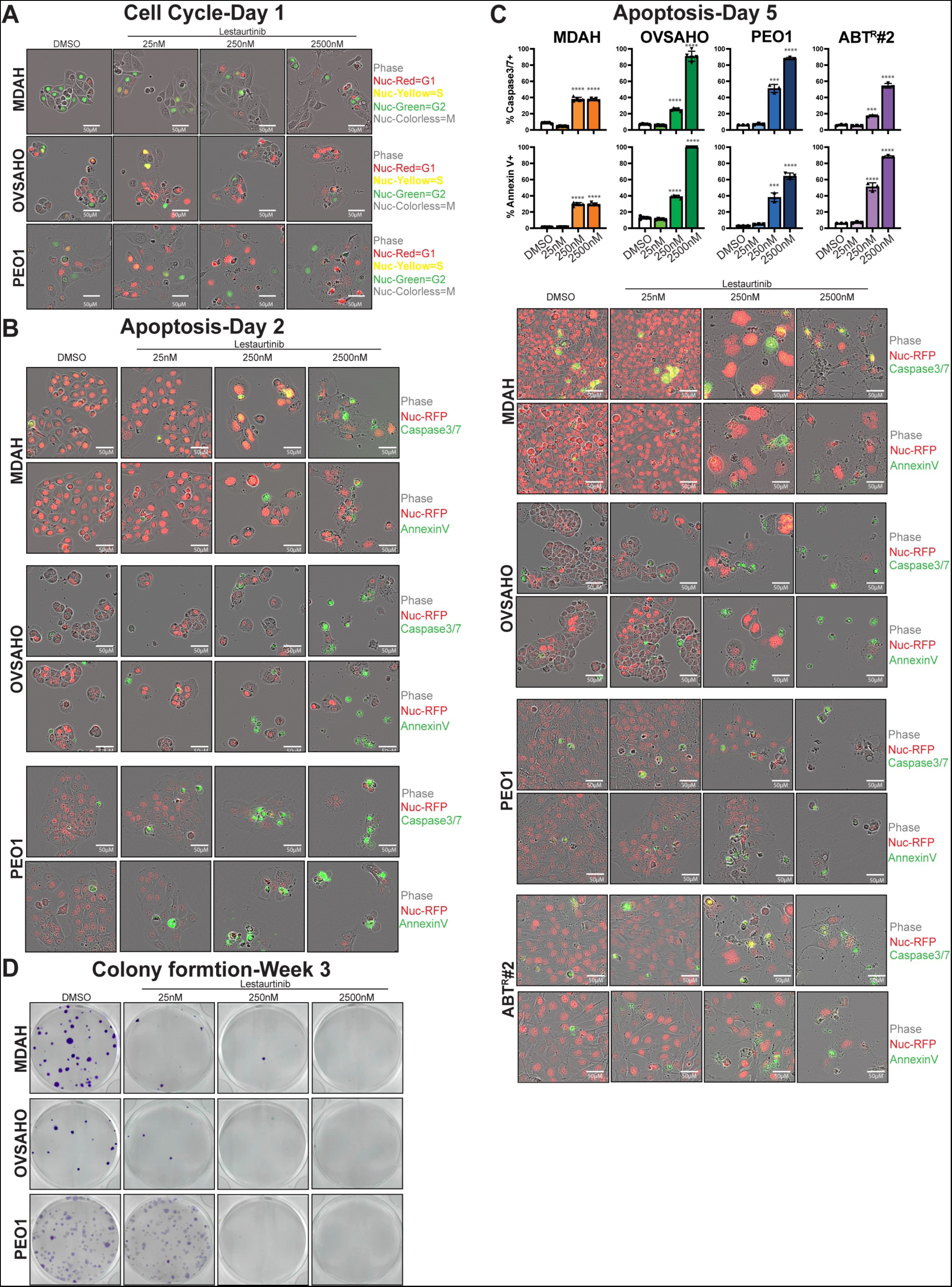
Lestaurtinib induces cell cycle arrest, apoptosis and prevents colony formation. **A)** Representative images of MDAH, OVSAHO, and PEO1-cell cycle reporter cells treated with indicated concentrations of lestaurtinib for 24 hours. **B)** Representative images of Nuc Red expressing cells treated with indicated apoptosis reporters and lestaurtinib for 2 days. **C)** Quantification of apoptosis at the 5-day timepoint. Percent apoptotic cells were defined as # of green positive cells to # of red positive cells. **D)** Representative images and quantification of colony formation 3 weeks after treatment with indicated concentrations of lestaurtinib. Abbreviations: Nuc = nuclear. Graphs represent mean ± standard error. ANOVA p-values: *<0.0332,**<0.0021,***<0.0002,****<0.0001.

**Supplemental Figure 3.**
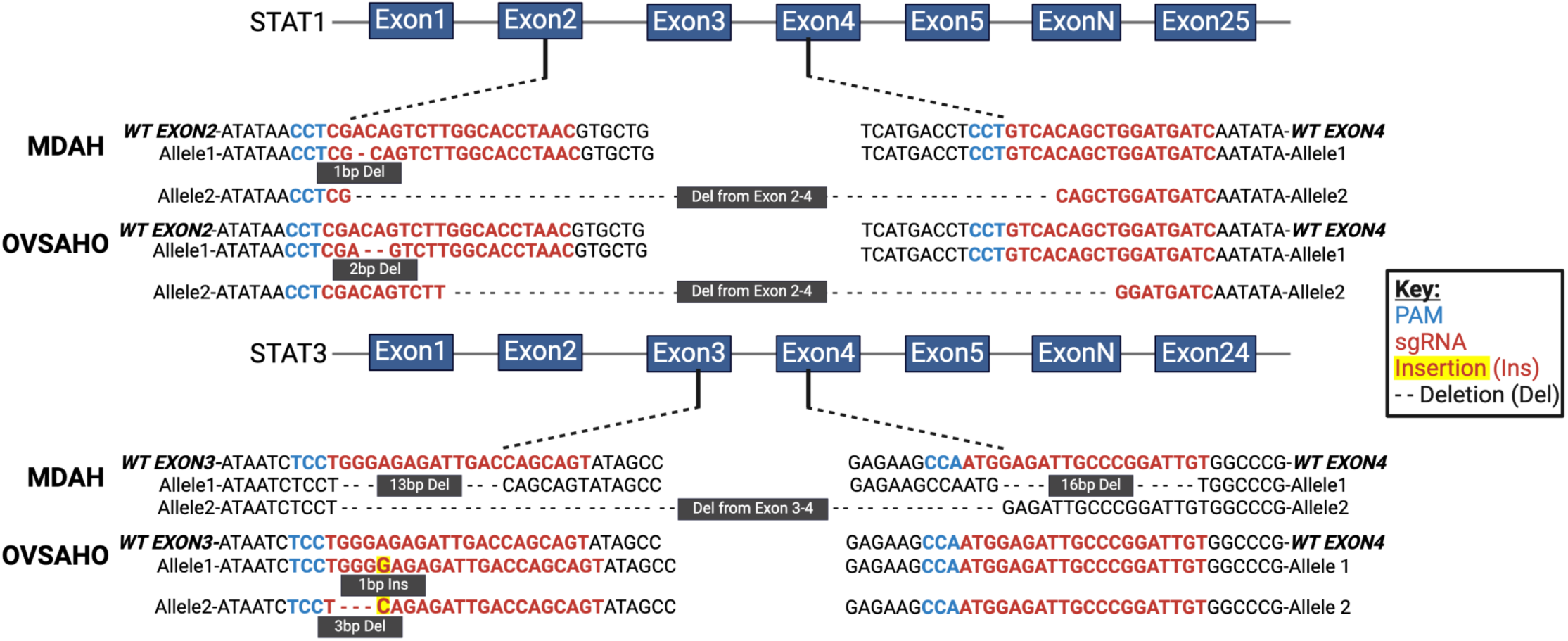
DNA sequencing confirms successful CRISPR KO of STAT1 and STAT3. DNA sanger sequencing results of STAT1 and STAT3 alleles in MDAH and OVSAHO STAT1 and STAT3 KO cells compared to the wildtype sequence. The sequence targeted by the guide RNA (sgRNA) is depicted in red, with the PAM sequence in blue. Genetic alterations in each allele are indicated. Abbreviations: WT = wildtype, Del = deletion, Ins = insertion. Figure created with BioRender.com.

**Supplemental Figure 4.**
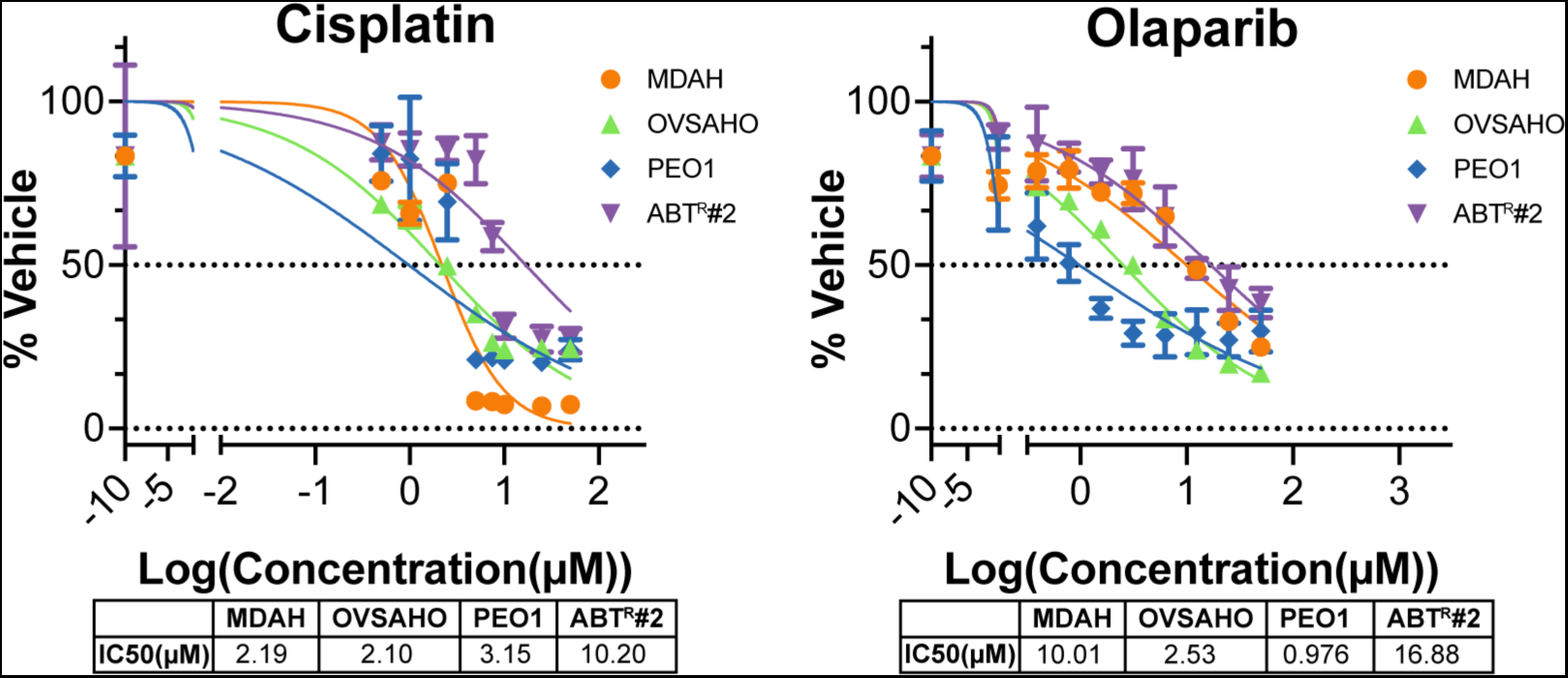
Indicated cell lines were treated with cisplatin (50, 25, 10, 7.5, 5, 2.5, 1, 0.5 µM) or olaparib (2-fold dilution starting at 50 µM) for 5-10 days and cell growth/viability was measured using crystal violet assays. IC50s were calculated using GraphPad Prism.

**Supplemental Figure 5.**
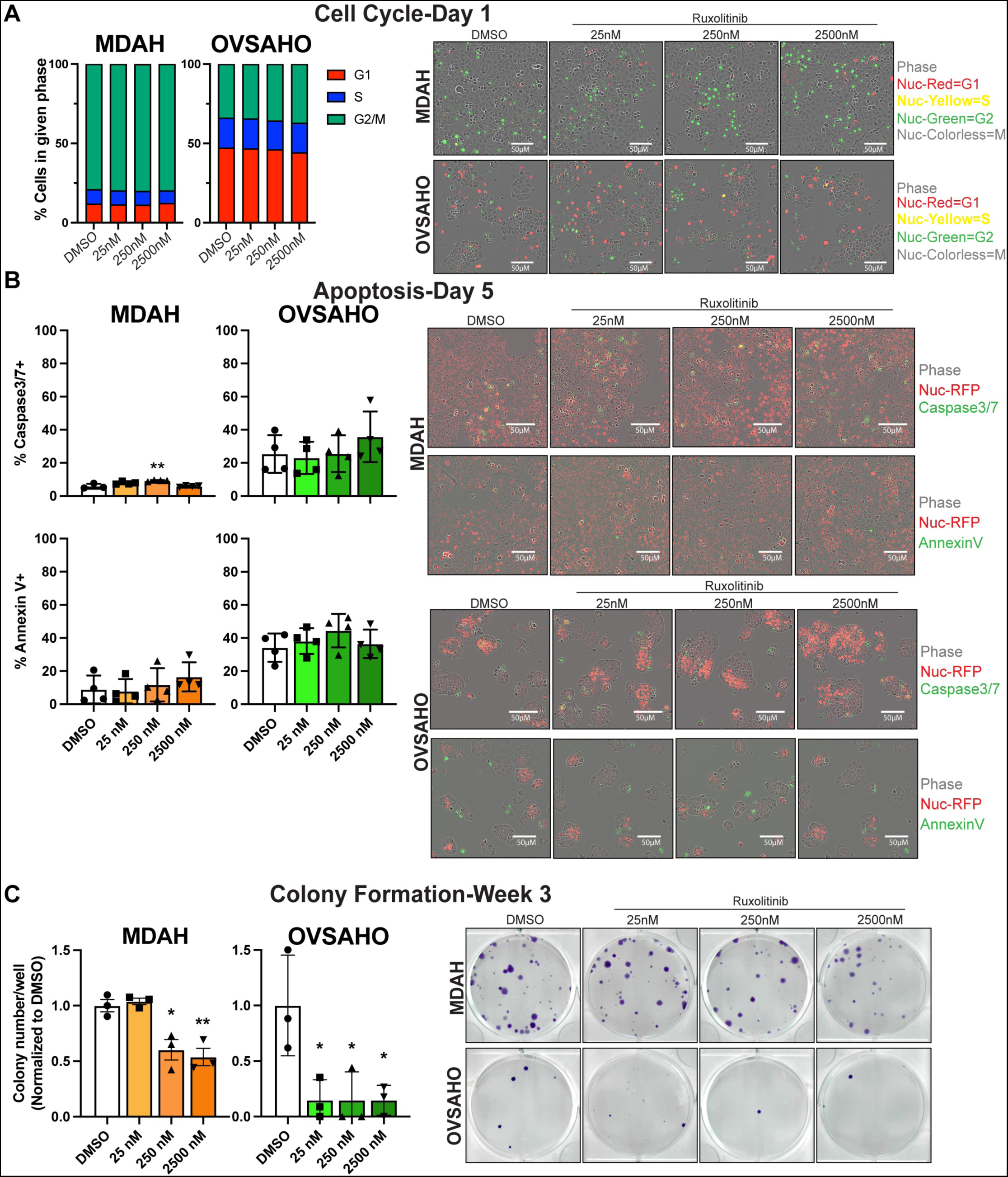
**A)** MDAH and OVSAHO-cell cycle reporter lines were treated with indicated concentrations of ruxolitinib for 24 hours and cell cycle phase was monitored. Percentage of cells in each phase was quantified via the IncuCyte S3 system based on nuclear red (G1), nuclear green (G2/M), and nuclear yellow (S) fluorescence. Representative images following treatment with ruxolitinib are shown. **B)** MDAH and OVSAHO-Nuc Red cell lines treated with Caspase 3/7-GFP or Annexin V-GFP reporter and indicated concentrations of ruxolitinib for 5 days. Percent apoptotic cells were quantified and defined as # of green positive cells to # of red positive cells. Representative images following 5 days of treatment are shown. **C)** Colony formation assays depicting the effects of ruxolitinib after 3 weeks of treatment. Representative images at the 3-week timepoint are shown. Abbreviations: Nuc = nuclear. Graphs depict mean ± standard error. ANOVA p-values: *<0.0332,**<0.0021,***<0.0002,****<0.0001.

**Supplemental Figure 6.**
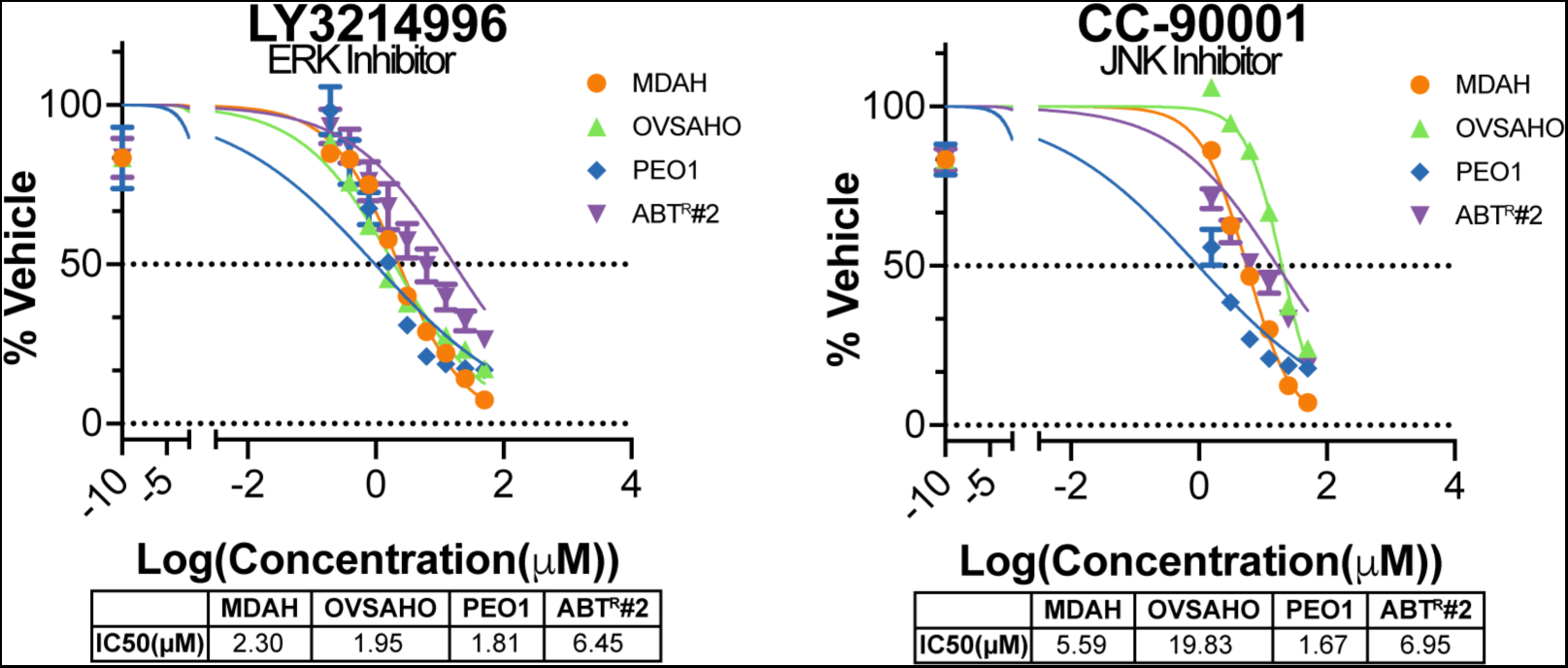
Indicated cell lines were treated with LY3214996 or CC-90001 (2-fold dilution starting at 50 µM) for 5-10 days and cell growth/viability was measured using crystal violet assays and IC50s were calculated using GraphPad Prism.

